# Comparative analysis of immunomodulatory effects of Artemisinin and Parthenolide on activation of immune cells and their roles in *Salmonella* Typhimurium infection

**DOI:** 10.64898/2026.09.12.751204

**Authors:** Tanisha Kumar, Aagosh Kishor Karhale, Shreyasee Das, Joel P. Joseph, Dipankar Nandi

## Abstract

Plant-derived immunomodulators are of immense therapeutic interest, of which sesquiterpene lactones (SLs) are a promising source. Artemisinin and Parthenolide are two structurally related SLs that are known to inhibit NF-κB. However, whether this mechanism produces comparable immunomodulatory outcomes has not been directly examined. In this study, we compared the effects of Artemisinin and Parthenolide across different *in vitro* and *in vivo* inflammatory contexts: T-cell activation, macrophage activation, and *Salmonella* Typhimurium infection. Parthenolide inhibited mouse T-cell activation more effectively than Artemisinin in terms of cell cycling, IL-2 production, and induction of activation markers, CD69 and CD44. In addition, Parthenolide suppressed production of LPS-induced nitrite and pro-inflammatory cytokines in primary thioglycolate (TG)-elicited macrophages as well as RAW 264.7 cells. It also reduced ROS production across all tested conditions. In contrast, Artemisinin exhibited comparatively modest effects, although it increased IL-6 in LPS-activated RAW 264.7 cells. Neither compound reduced bacterial burden in an *in vitro* model of *S.* Typhimurium infection; however, Parthenolide, but not Artemisinin, lowered TNF-α amounts. Together, these findings demonstrate that Parthenolide is a broader and more consistent immunomodulator than Artemisinin. These observations led us to investigate the effects of Parthenolide in an *in vivo* model of mice orally fed with *S*. Typhimurium. Parthenolide-treated mice showed higher survival accompanied with lower sera amounts of IL-6 and TNF-α, indicating that its protective effects operate by limiting host immunopathology. The implications of these observations on the use of compounds that lower host inflammatory responses during infections are discussed.

## INTRODUCTION

The immune system operates as a finely tuned system of checks and balances, requiring both the capacity to mount a defense against pathogens and restraint to avoid tissue damage. Compounds that can shift this balance, either toward greater activation or greater restraint, are broadly termed immunomodulators. They can be further classified as immunostimulants or immunosuppressants depending on the direction of their effects (1,2). Immunostimulants, such as interferons and interleukin-based therapies, enhance immune function by activating dendritic cells, natural killer cells, or T-cell populations, and are used against malignancies and chronic viral infections such as hepatitis C (1). Immunosuppressants, including calcineurin inhibitors like cyclosporine and TNF-α-targeting biologics, act in the opposite direction, inhibiting T-cell proliferation or neutralizing pro-inflammatory mediators to manage autoimmune conditions such as rheumatoid arthritis and inflammatory bowel disease (2). Both classes remain constrained by a persistent problem in drug development, that synthetic compounds with sufficient selectivity and potency are difficult to achieve without introducing toxicity. This has driven growing interest in plant-derived alternatives, because many naturally occurring phytocompounds regulate immune activity through mechanisms distinct from, and often better tolerated than, their synthetic counterparts (3).

Sesquiterpene lactones (SLs), a class of sesquiterpenoids defined by a lactone ring and concentrated largely in the Asteraceae family, are reported to have anti-inflammatory and immunoregulatory activity acting through NF-κB, MAPK, and JAK-STAT signaling across a range of cellular and *in vivo* model systems (4,5). A previous study, from our laboratory reported one such compound, 7-Hydroxy Frullanolide, derived from methanolic extract of *Sphaeranthus indicus*. It was shown to inhibit CD4^+^ T cell and peritoneal macrophage activation and ameliorate DSS-induced colitis in mice by increasing intracellular calcium amounts (6). Building on this earlier work, we sought to test whether other well-characterized SLs, would produce comparable immunomodulatory outcomes.

Artemisinin, isolated from *Artemisia annua*, is best known as an antimalarial discovered by Tu Youyou (7), but it also modulates a broad range of immune cells, including neutrophils, dendritic cells, macrophages, and T-cell subsets (8). Parthenolide, derived from feverfew (*Tanacetum parthenium*), inhibits NF-κB more directly, by blocking IκB kinase activity and p65 DNA binding, and has additionally been reported to covalently engage JAK2 (5) and the protein Trim33, stabilizing Smad4 to suppress NF-κB signaling and improve outcomes in a mouse model of sepsis (9). Although both compounds converge on NF-κB inhibition (10,11), whether this shared mechanism produces comparable effects on immune cell activation, cytokine production, and redox homeostasis remains untested under matched experimental conditions. This aspect is important as existing reports derive from disparate cell types and disease models that preclude direct comparison. In this study, we compared the effects of Artemisinin and Parthenolide across different inflammatory mouse model systems: T-cell activation, macrophage activation — a tumor cell line and primary TG-elicited peritoneal macrophages, *in vitro* and *in vivo* infection of *Salmonella* Typhimurium. This comparative study demonstrates the ability of Parthenolide to lower inflammatory responses *in vitro* as well as *in vivo* which may be useful in lowering host-mediated inflammatory responses during infections and diseases.

## MATERIALS AND METHODS

### Cell culture

RAW 264.7 (monocyte/macrophage) cell line was cultured in DMEM with high glucose, supplemented with 320 μg/mL glutamine, along with 100 μg/mL penicillin, 250 μg/mL streptomycin, 50 μg/mL Gentamicin, 3.5 μL/L β-mercaptoethanol and 10% FBS. The cells were maintained at 37°C in a humidified incubator with 5% CO_2_ and tested to be free of Mycoplasma.

### Animals

All experiments were performed on 6-8 weeks old male C57BL/6 mice. All experiments were conducted as per the Control and Supervision rules 1998 of the Ministry of Environments and Forests Act, Government of India, and the Institutional Animal Ethics Committee of the Indian Institute of Science, under the permit number: CAF/Ethics/976/2023. Mice were bred and maintained in the Central Animal Facility, IISc, Bengaluru. Animals were housed in standard mouse cages under conditions of optimum light (12:12 h light-dark cycle), temperature (22 ± 1°C), and humidity (50-60%), with access to food and water *ad libitum*.

### Isolation of CD3^+^ T cells

The CD3^+^ T cells isolated from the inguinal and mesenteric lymph nodes of 6-8 weeks old male C57BL/6 mice were used for all the *in vitro* experiments. Lymph nodes were disrupted mechanically and passed through a cell strainer to obtain a single cell suspension. The suspension was incubated in a T25 flask coated with 100 μg/mL of AffiniPure goat anti-mouse IgG antibody (Jackson Immuno Research laboratories, USA) for 30 min in a humidified incubator at 37°C twice, to deplete B cells (12). The purity of CD3^+^ T cells was estimated to be more than 97% positive using flow cytometry (Supplementary Figure 1).

### Activation of CD3^+^ T cells

CD3^+^ T cells were seeded in 96-well U-bottom plates at a density of ∼50,000 cells/well in RPMI media supplemented with 5 % FBS and activated using PMA (10 ng/mL) and Ionomycin (0.1 µM), a TCR independent activation model. For compound treatment, the cells were pretreated with the specific concentrations of Artemisinin or Parthenolide at 37°C for 30 min before activating with PMA and Ionomycin. Cell-free culture supernatants were collected for cytokine measurements, while the cells were collected and fixed for cell cycle analysis or ROS estimation at 36 h post activation (12,13).

### Isolation of TG-elicited peritoneal macrophages

C57BL/6 mice (20–25 g) were subjected to intraperitoneal injection of 1 mL 4% Brewer’s TG broth (Sigma Aldrich, USA), which was prepared, autoclaved, and aged for at least one month before use. Four days post-injection, the mice were sacrificed and peritoneal exudate cells (PECs) were harvested by flushing the peritoneal cavity with 1x ice cold PBS. Cells were resuspended in DMEM medium supplemented with 10% FBS and plated at a density of approximately 0.15 × 10 cells per well in a 96-well flat-bottom plate. Non-adherent cells were removed 1 h after seeding. The adherent peritoneal macrophages (6,14) were then utilized for subsequent experiments (Supplementary Figure 3A).

### Activation of macrophages

The peritoneal macrophage cells were activated with 1 µg/mL of lipopolysaccharide (LPS) from *E. coli* O111:B4 (Sigma Aldrich, USA). Cells were pre-treated with the specific concentrations of Artemisinin or Parthenolide for 30 min. RAW 264.7 cells were activated with 1µg/mL LPS or 25 U/mL IFN-γ alone, or in combination, post treatment with Artemisinin or Parthenolide at two doses (1 µM and 5 µM) for 30 min. After 36 h of incubation the cell-free culture supernatants were used to estimate nitrite and cytokine levels, while the cells were collected for ROS estimation (6,14).

### Nitrite Measurements

Nitric oxide production by macrophage was quantified by measuring nitrite amounts in cell-free supernatants using the Griess assay (14).The Griess reagent was prepared using 1% (w/v) sulfanilamide and 0.1% (w/v) N-(1-naphthyl) ethylenediamine dihydrochloride, which were dissolved in a 2.5% ortho-phosphoric acid solution prepared in MilliQ water. In the assay condition, the reaction mixture was prepared by combining 25 μL of cell-free supernatant, 25 μL of MilliQ water, and 100 μL of Griess reagent. The blank control consisted of 25 μL of DMEM medium fortified with 5% FBS, 25 μL of MilliQ water, and 100 μL of Griess reagent. Absorbance at 550 nm was measured using a micro-plate reader (Infinite 200Pro, TECAN, Switzerland), and nitrite concentrations in supernatants were calculated from a sodium nitrite standard curve in the range from 0.39 to 100 μM.

### Cytokine Measurements

Amounts of IL-2, IL-6, and TNF-α in cell-free culture supernatants were quantified using the respective eBioscience™ ELISA Ready-SET-Go™ kit, following the manufacturer’s protocol. The standard curves for the respective cytokines were generated by measuring absorbance at serial dilutions ranging from 31.25 to 2000 pg/mL. Cytokine concentrations in the samples were determined by interpolation from their respective standard curves within the assay’s detection range (12).

### Alamar Blue Staining

Post 36 h of activation and treatment, AlamarBlue HS cell viability reagent (Invitrogen) was added directly to each well at a final concentration of 10% and plates were incubated for 2–4 h at 37°C in a 5% CO atmosphere, protected from light. Fluorescence intensity was measured using a plate reader at an excitation wavelength of 560 nm and emission wavelength of 590 nm.

### Flow Cytometry

For cell cycle analysis, T cells were collected post 36 h of activation and fixed with 70% ethanol on ice for at least 4 h. The cells were then washed with 1x PBS and treated with 100 μg/mL of RNase A (Qiagen, USA) followed by staining with 50 μg/mL of propidium iodide (Sigma-Aldrich, USA). The cells were incubated in the dark at room temperature for 10-15 min and acquired on the flow cytometer (BD FACS Verse, BD Biosciences, USA) (12). To estimate activation of T cell markers, the cells were stained for surface markers by incubating with a cocktail of pre-titrated dilutions (anti-mouse CD3 at 1:200, anti-mouse CD4 at 1:200, anti-mouse CD69 at 1:200 and anti-mouse CD44 at 1:200) of the conjugated antibodies and fixable cell viability dye Zombie NIR Fixable Viability dye (BioLegend, USA) at 1:1000 (12). The cells were incubated in the dark on ice for 45 min with intermittent tapping. The cells were washed with FACS Buffer (PBS + 2% FBS + 0.1% Sodium Azide) and fixed with 1% paraformaldehyde in PBS. Data were acquired on Cytoflex^LX^ flow cytometer (Beckman Coulter Life Sciences, Indianapolis, IN, USA). For ROS estimation, post 36 h of activation and treatment, T cells and peritoneal macrophages were collected, washed and stained with 2,7- dichlorofluorescein diacetate (DCF-DA, Sigma-Aldrich, USA) for 45 min at 37°C. The fluorescence intensity was measured using flow cytometry. Data from the flow cytometer were analysed using the FlowJo software. Events were gated based on FSC-A versus SSC-A and histograms were generated. The percentage of positive cells and the median fluorescence intensity were estimated by gating the desired populations.

### *In vitro Salmonella* Typhimurium infection

Approximately 24 h before initiating the experiment, RAW 264.7 cells were seeded at the density of 1.2×10^4^ cells/well in a 96-well plate. *Salmonella* Typhimurium were grown overnight at 37°C. The absorbance of the overnight grown pre-inoculum was normalized to OD 2.0 at 600 nm and 50 μL of this pre-inoculum was inoculated in 50 mL of LB and grown at 37°C for 10 h. RAW 264.7 cells were infected with an MOI 1:10 following incubation at 37°C for 60 min. The cells were washed with PBS and 100 μL of 100 μg/mL of Gentamicin in DMEM was added to kill the extracellular bacteria (15). To study effects of compounds on intracellular replication, Gentamicin-containing medium was discarded, and cells were incubated with Artemisinin or Parthenolide diluted in antibiotic free media at 37°C. At 2 and 18 h post infection, supernatants were collected for cytokine estimation, and the cells were washed twice with PBS, followed by lysis with 0.1% Triton X-100. Appropriate dilutions were plated on Salmonella-Shigella (SS) agar plates to determine the intracellular bacterial load (15).

### *In vivo* infection in mice via oral feeding of *Salmonella* Typhimurium

Six- to eight-week-old, male, C57BL/6 mice (n= 3-4 mice per group) were orally infected with ∼ 1×10^6^ CFU/mouse of *S*. Typhimurium. Post 2 days of infection Parthenolide was injected intraperitoneally (i.p.) at dose of 10mg/kg or DMSO as vehicle control group. As a control, mice were injected with 1.25 mL/kg of DMSO. Post 4 days of infection, mice were sacrificed, organs were harvested, weighed and homogenised to determine the bacterial count. Blood was collected from retro-orbital sinus to determine the cytokine levels from isolated sera (15). For the survival experiments, C57BL/6 mice were infected with *S.* Typhimurium through the oral route. Subsequently, the mice were treated intraperitoneally 2 days post infection with Parthenolide at a dose of 10mg/kg and survival of the mice was monitored over 14 days.

### Statistical Analysis

All graphical representations and statistical analyses were performed on GraphPad Prism software version 9.5.1. All data are presented as mean ± standard error of the mean (SEM). Statistical analyses were conducted using one-way ANOVA with Tukey’s multiple comparisons test with single pooled variance or two-way ANOVA with Tukey’s multiple comparisons test. A *p*-value of less than 0.05 was considered statistically significant, with significance levels denoted as \**p* < 0.05, \*\**p* < 0.01, \*\*\**p* < 0.001, and \*\*\*\**p* < 0.0001 for comparisons between conditions.

## RESULTS

### Parthenolide inhibits T cell activation more efficiently than Artemisinin

To elucidate the effects of Artemisinin and Parthenolide (Figure 1 A, B) on T cell activation and proliferation, we pretreated primary T cells isolated from C57BL/6 mice (Supplementary Figure 1) with different concentrations of compounds (1 µM, 5 µM), followed by activation using PMA and Ionomycin after 30 min (12,13). Activation with PMA and Ionomycin resulted in formation of blast zones, and treatment with Artemisinin showed no significant decrease in the blast zones compared to the activated cells. However, upon treatment with Parthenolide, a dose dependent decrease in blast zones could be seen (Figure 1 C). Furthermore, IL-2 levels in the cell-free supernatant showed no difference upon treatment with Artemisinin but decreased upon Parthenolide treatment (Figure 1 D). To support these observations, we investigated the changes in activation associated proliferation of T cells upon treatment with the compounds. The percentage of cycling and hypodiploid cells were determined using flow cytometry. A significant decrease in the percent of cycling cells and an increase in the percentage of hypodiploid cells was observed upon treatment with Parthenolide. No such changes were observed when the cells were treated with Artemisinin (Figure 2 A, B). This is suggestive that Parthenolide is better at inhibiting T cell-activation.

**FIGURE 1.**
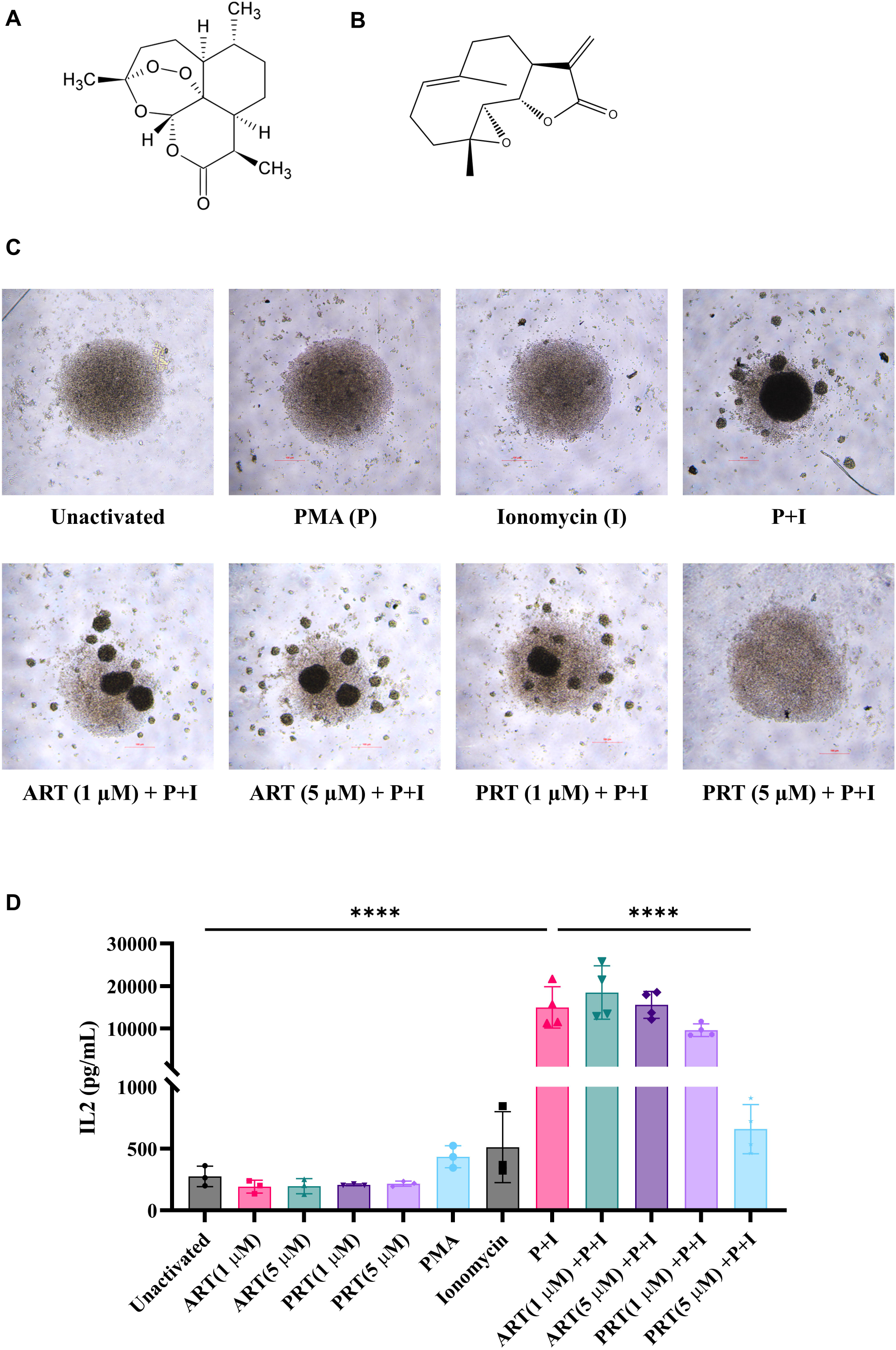
Parthenolide inhibits T cell activation-associated changes in morphology and IL-2 production more effectively than Artemisinin. Molecular structure of (A) Artemisinin and (B) Parthenolide. (C) Brightfield images of T cells activated with 10 ng/mL PMA and 0.1 μM Ionomycin and treated with different concentrations of Artemisinin and Parthenolide. (D) IL-2 levels of T cells activated with PMA and Ionomycin and treated with different concentrations of Artemisinin and Parthenolide. Data are represented as mean ± SEM from three independent experiments (n=3). One way ANOVA was performed to test the statistical significance, where ****p<0.0001.

**FIGURE 2.**
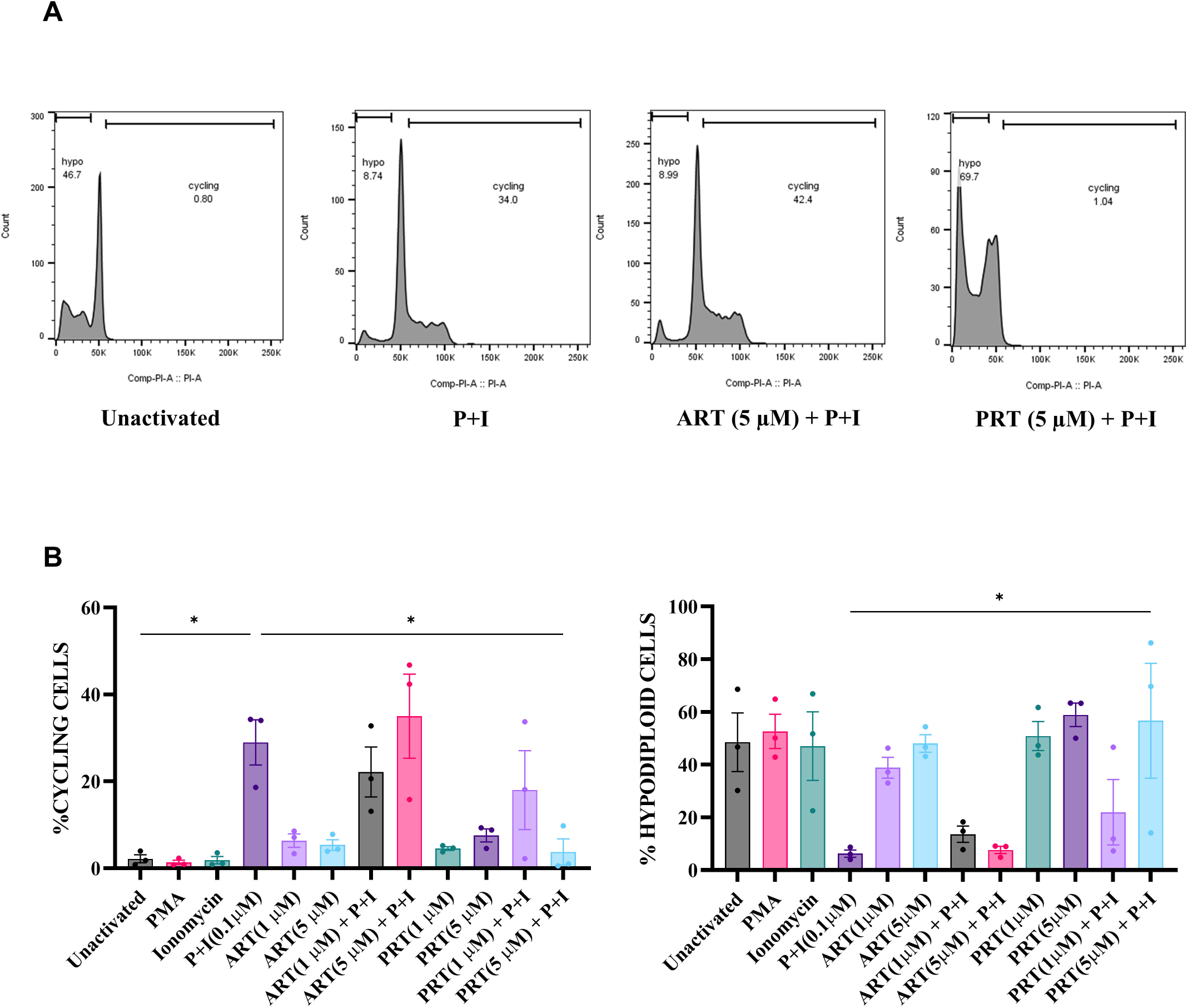
Parthenolide inhibits T cell activation-associated changes in cell cycling more effectively than Artemisinin. (A) Representative flow cytometry plots of cell cycling and hypodiploid T cells activated with PMA and Ionomycin and treated with 5 μM of Artemisinin and Parthenolide. (B) Percentage of cycling and hypodiploid cells quantified from the flow cytometric plots. Data are represented as mean ± SEM from four independent experiments (n=4). One-way ANOVA was performed to test the statistical significance, where *p<0.05, **p<0.01.

### Parthenolide affects expression of early and late activation markers of T cells

To further investigate the effects of Artemisinin and Parthenolide on T cell activation, we evaluated the expression of T cell activation markers CD69 and CD44 using flow cytometry. A time kinetics experiment done previously showed that T cells expressing CD69 increased at 12 h post activation and T cells expressing CD44 increased at 36 h post activation (12). We did observe a significant increase in T cells expressing CD69 and CD44 upon activation with PMA and Ionomycin. Pretreatment with Parthenolide at 5 μM resulted in a significant decrease in the T cells expressing CD69 and CD44. However, there was no decrease in CD69 or CD44 expression upon Artemisinin treatment (Figure 3, Supplementary Figure 2). This correlates with our previous observation that Parthenolide is better at inhibiting T cell activation compared to Artemisinin.

**FIGURE 3.**
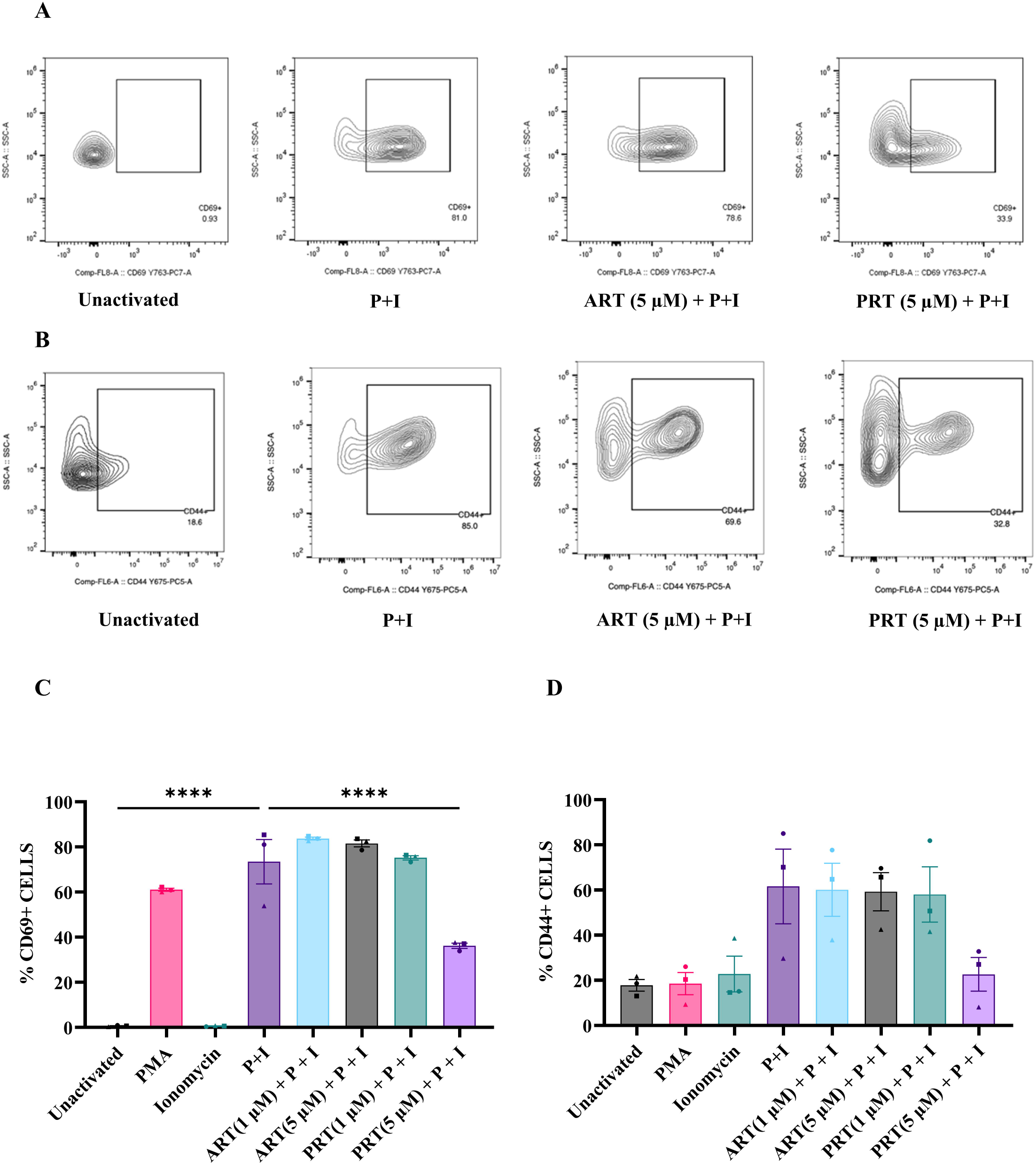
Parthenolide treatment lowers expression of T cell surface activation markers. (A) Representative flow cytometry plots of CD69 surface expression on T cells. T cells were treated with 5 μM of Artemisinin or Parthenolide prior to activation with PMA and Ionomycin and were assessed for CD69 expression post 12 h. (B) Representative flow cytometry plots of CD44 surface expression on T cells. T cells were treated with 5 μM of Artemisinin or Parthenolide prior to activation with PMA and Ionomycin and CD44 expression was assessed post 36 hours. (C) Percentage of CD69^+^ T cells at 12 h post activation. (D) Percentage of T cells positive for CD44 expression at 36 h post activation. Data are represented as mean ± SEM from three independent experiments (n = 3). One-way ANOVA was performed to test the statistical significance, where ****p<0.0001.

### Parthenolide reduces ROS amounts and metabolic activity in activated T cells

In mouse T cells, activation with PMA and Ionomycin increased the total intracellular ROS amounts, as evidenced by the enhanced fluorescence of the ROS-sensitive DCF-DA dye (12).

Although Artemisinin had no effect on intracellular ROS production initially at 12 h, it further increased activation-induced ROS generation at 36 h. On the contrary, treatment with Parthenolide decresed the intracellular ROS levels in activated T cells even at an earlier timepoint (12 h) (Figure 4 A, B).

**FIGURE 4.**
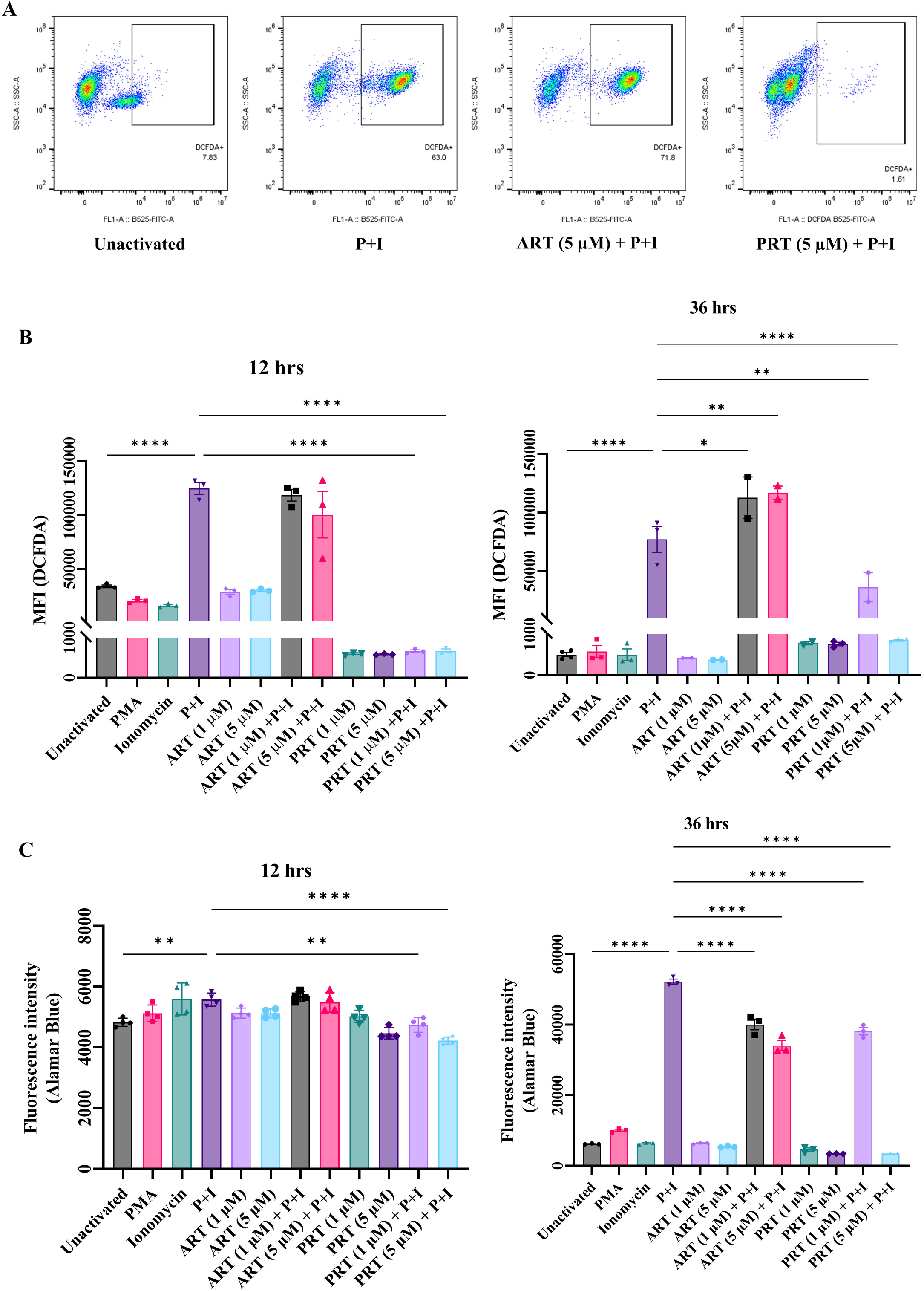
Parthenolide reduces intracellular ROS amounts and metabolic activity in activated T cells. (A) Representative flow plots of DCF-DA staining of unactivated T cells, T cells activated with PMA and Ionomycin and treated with different doses of Artemisinin and Parthenolide at 36 h post activation. (B) Mean fluorescence intensity of DCF-DA staining of T cells activated and treated with different doses of Artemisinin and Parthenolide 12 h and 36 h post activation. (C) Fluorescence intensity of Alamar Blue staining of activated T cells treated with different doses of Artemisinin or Parthenolide post 12 or 36 h of activation. Data are represented as mean ± SEM from 3-4 independent experiments (n = 3). One way ANOVA was performed to test the statistical significance, where *p<0.05, **p<0.01, ****p<0.0001.

Overall metabolic activity, assessed by Alamar Blue assay, increased significantly upon activation with PMA and Ionomycin. Both Artemisinin and Parthenolide treatments decreased the metabolic activity of the cells; however, the decrease upon Parthenolide treatment was more pronounced and robust, as seen even at an earlier timepoint (Figure 4 C). These results are suggestive of more potent inhibitory and antiproliferative effects of Parthenolide on activated T cells.

### Parthenolide inhibits LPS-induced pro-inflammatory responses of TG-elicited peritoneal macrophages

To understand the effect of Artemisinin and Parthenolide on other immune cells, we used TG-elicited peritoneal macrophages. The isolation procedure yielded ∼96% F4/80^+^ macrophages, as measured by flow cytometry (Supplementary Figure 3) and pretreated them with different doses of Artemisinin and Parthenolide followed by activation with LPS. Although neither compound induced any significant change to cell morphology (Supplementary Figure 4), both compounds induced differential amounts of nitric oxide and pro-inflammatory cytokine production. Parthenolide significantly inhibited LPS-induced nitrite levels, whereas Artemisinin only slightly decreased nitrite levels (Figure 5 A). Upon cytokine estimation, we observed that Parthenolide decreased LPS-induced IL6 production but had no effect on TNF-α production; however, Artemisinin treatment showed no difference (Figure 5 B, C). The cells showed no significant change in metabolic activity as measured by Alamar Blue assay, probably because these cells are non-proliferating (Figure 5 D). Subsequently, there was a decrease in LPS-induced ROS levels upon both Artemisinin and Parthenolide treatment, with Parthenolide showing a more significant effect (Figure 5 E, F). These results are suggestive of Parthenolide as a more potent anti-inflammatory compound.

**FIGURE 5.**
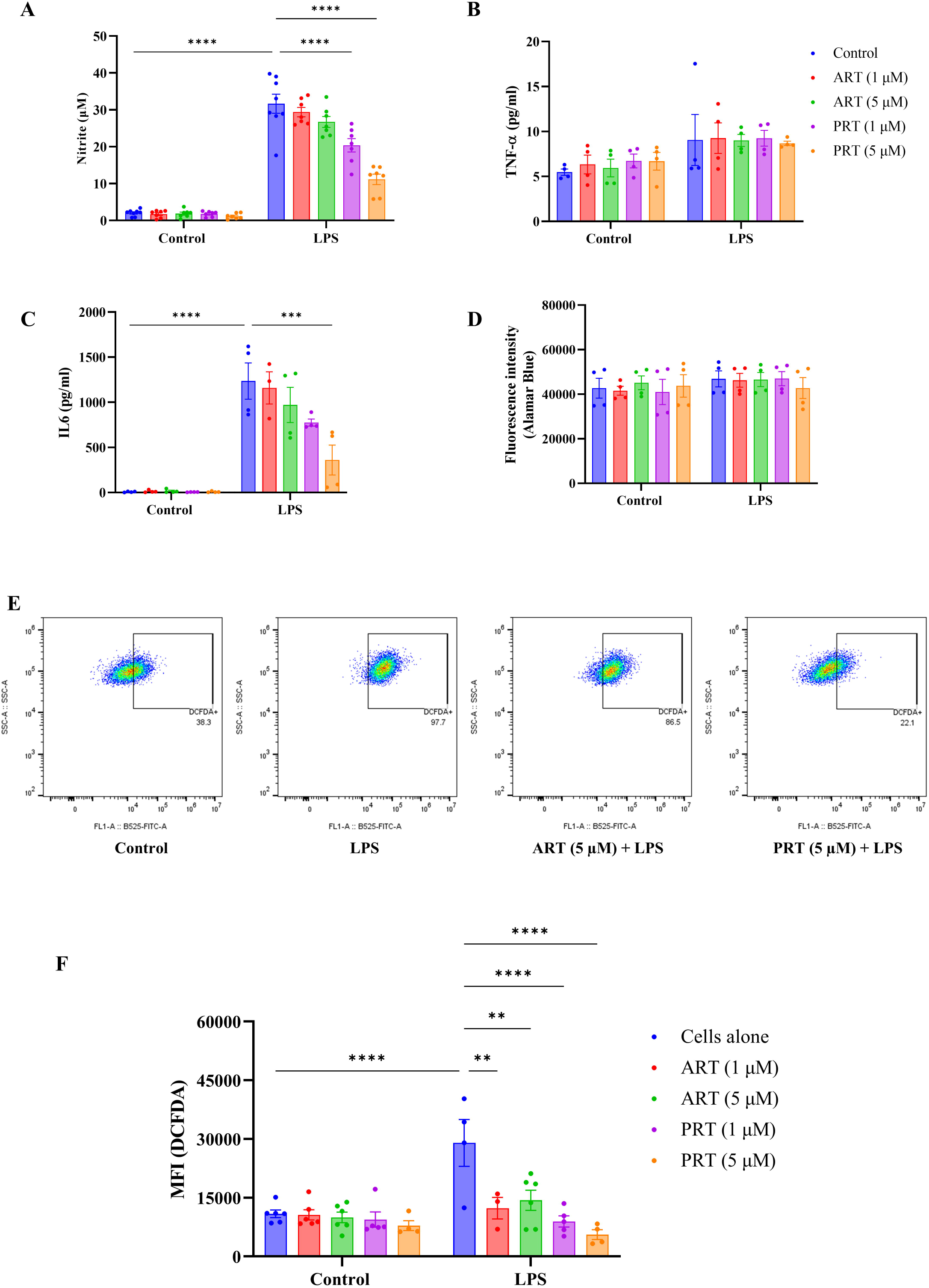
Parthenolide inhibited LPS-induced nitrite and IL6 responses as well as intracellular ROS levels in TG-elicited peritoneal macrophages. (A) Nitrite, (B) TNF-α and(C) IL-6 levels, in the cell free culture supernatants of TG-elicited peritoneal macrophages treated with different concentrations of Artemisinin and Parthenolide in the presence or absence of LPS at 36 h post activation. (D) Fluorescence intensity of Alamar Blue staining of LPS-activated adherent peritoneal macrophages and treated with different concentrations of Artemisinin and Parthenolide. (E) Representative flow plots of DCF-DA staining of TG-elicited peritoneal macrophages, activated with LPS and treated with 5 µM of Artemisinin and Parthenolide at 36 h post activation. (F) Mean fluorescence intensity of DCF-DA staining of LPS-activated adherent peritoneal macrophages and treated with different doses of Artemisinin and Parthenolide. Data are represented as mean ± SEM from 4-6 independent experiments. Two-way ANOVA was performed to test the statistical significance, where **p<0.01, ***p<0.001, ****p<0.0001.

### Parthenolide inhibits activation-induced responses in proliferating macrophages

Primary peritoneal macrophages are largely non-proliferative; therefore, we used the RAW 264.7 macrophage cell line to examine the *in vitro* effects of Artemisinin and Parthenolide on a proliferating tumor macrophage cell line under inflammatory conditions. The cells were activated with LPS and IFN-γ separately and in combination and treated with different doses of Artemisinin and Parthenolide. Treatment with the compounds resulted in decrease in activation-induced nitrite levels, which was more significant with Parthenolide (Figure 6 A). Upon cytokine estimation, we observed that Artemisinin elevated IL-6 beyond the levels observed with activation alone. Parthenolide exerted the opposite effect, reducing activation-induced IL-6 and TNF-α (Figure 6 B, C). Upon activation, RAW 264.7 cells showed decrease in overall metabolic activity as measured by Alamar Blue. While Parthenolide partially rescued the activation-induced decrease in metabolic activity, Artemisinin showed no significant effect (Figure 6 D). In summary, Parthenolide proved to be better at suppressing the inflammatory responses of LPS and IFN-γ induced inflammation in RAW 264.7 macrophages.

**FIGURE 6.**
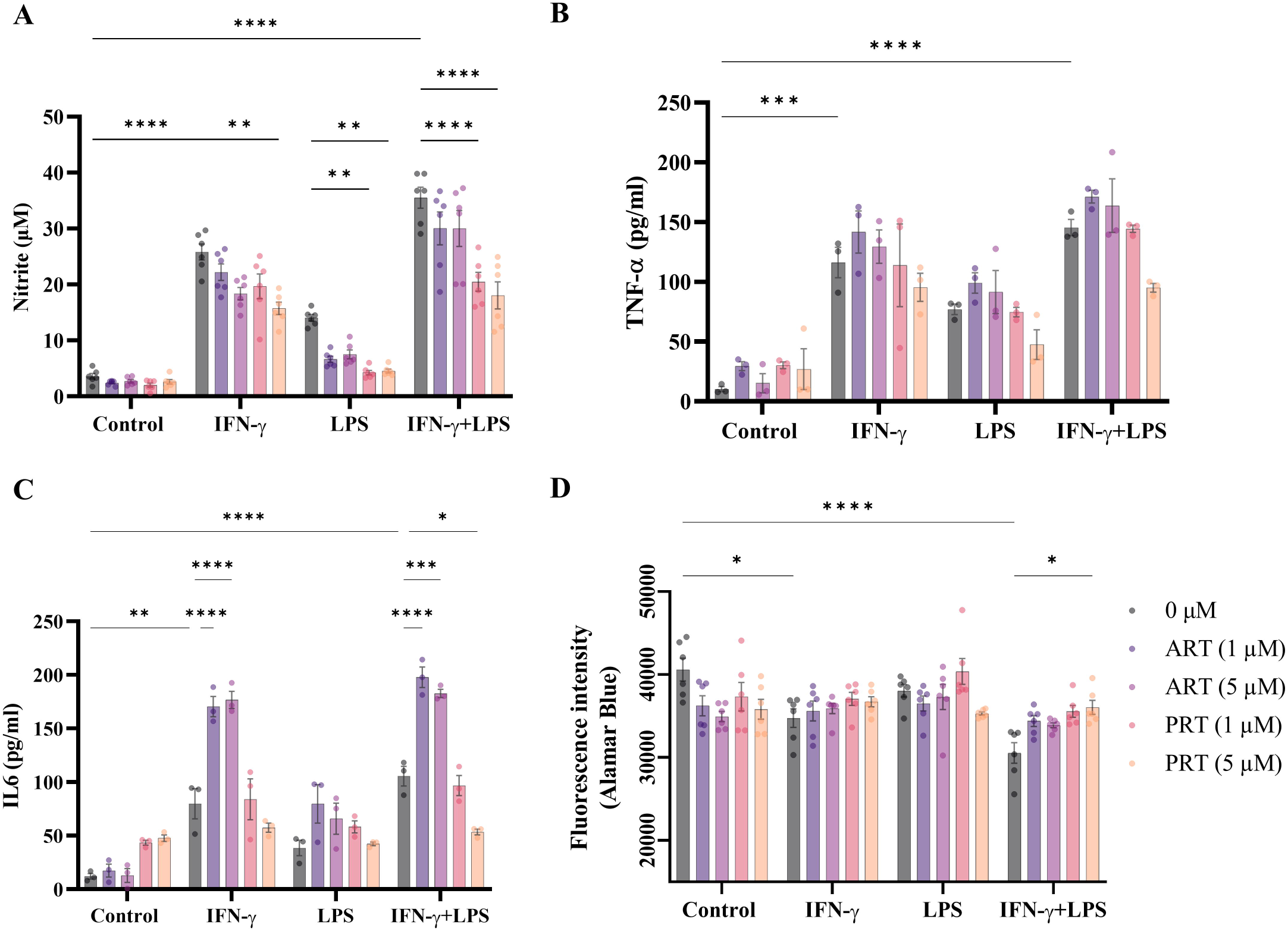
Parthenolide inhibits activation-induced responses in RAW 264.7 cells. (A) Nitrite, (B) TNF-α and (C) IL-6 levels, in the cell-free culture supernatants of RAW 264.7 cells treated with different concentrations of Artemisinin and Parthenolide in the presence or absence of LPS and IFN-γ, alone and in combination, at 36 h post activation. (D) Fluorescence intensity of Alamar Blue staining of activated and unactivated RAW 264.7 cells and treated with different concentrations of Artemisinin and Parthenolide. Data are represented as mean ± SEM from five independent experiments (n = 5). Two-way ANOVA was performed to test the statistical significance, where *p<0.05, **p<0.01, \*\*\**p*<0.001, ****p<0.0001.

### Parthenolide decreases infection-induced TNF-α production in RAW 264.7 macrophages upon *in vitro* infection

From the above observations, Parthenolide appeared to be a more effective immunomodulatory compound. To further test the effects of these compounds in an infection model, we first evaluated the compounds for their direct bacteriostatic effects and found that both compounds could not reduce bacterial growth at the tested concentrations (Supplementary Figure 5). We further examined whether treatment with the compounds could contribute to a reduction in the bacterial burden within immune cells. RAW 264.7 cells were infected with *S.* Typhimurium at a MOI of 1:10 and intracellular bacterial replication was studied at 2 h and 18 h post infection (Figure 7 A). Bacterial burden, measured as log CFU/mL, increased over this period. Treatment with Artemisinin or Parthenolide however, did not affect the number of viable intracellular bacteria (Figure 7 B). We further investigated the levels of IL-6 and TNF-α in the cell-free supernatant to assess whether infection-driven cytokine responses were affected by treatment. Post 18 h of infection, there was a significant increase in the TNF-α levels, which reduced upon Parthenolide treatment (Figure 7 C, D). This suggests that Parthenolide may not directly decrease bacterial burden, but it can lower the inflammatory responses by immune cells during infection.

**FIGURE 7.**
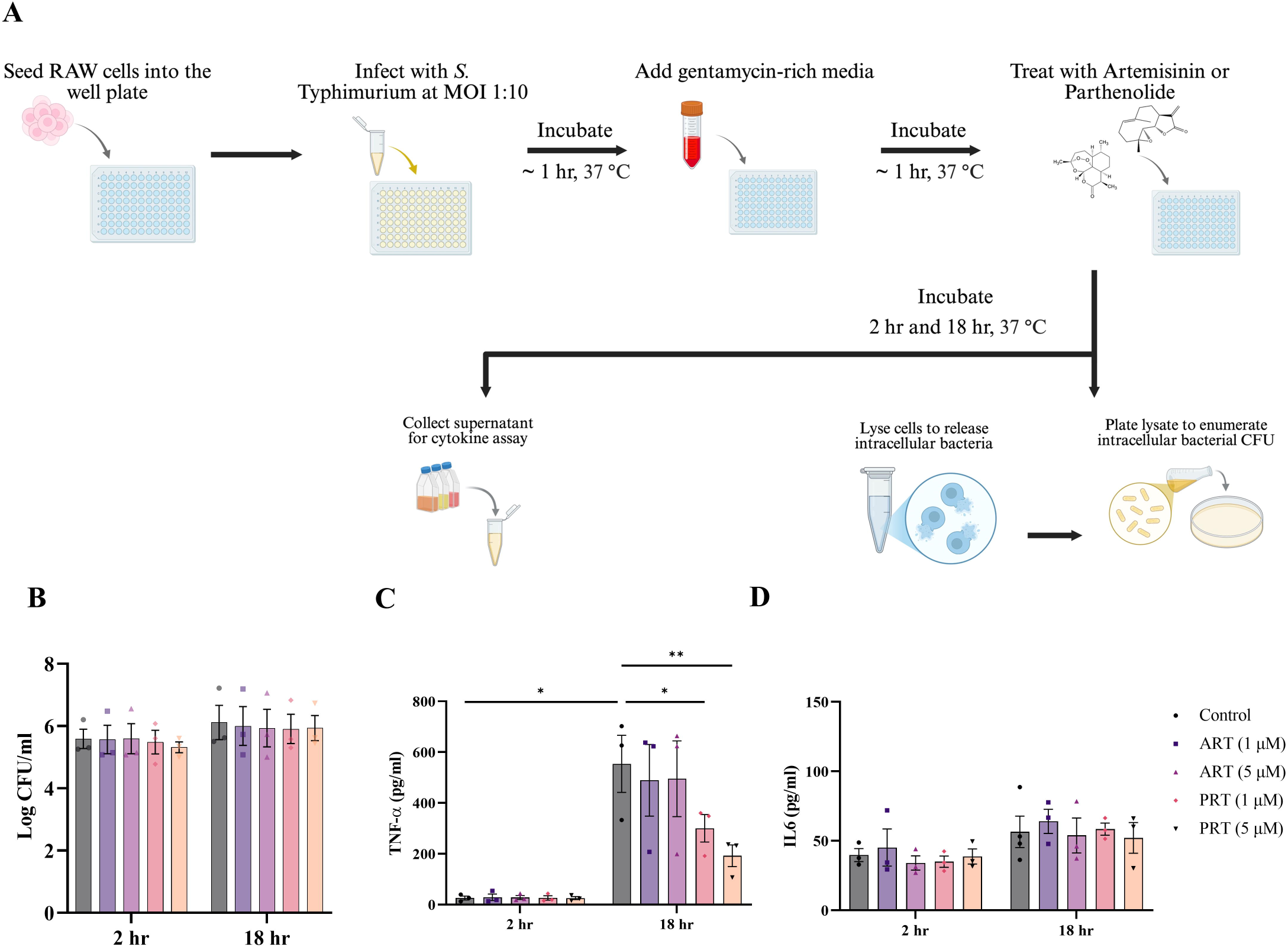
Parthenolide lowers *S*. Typhimurium induced TNF-α production in an *in vitro* infection model in RAW 264.7 cells. (A) Experimental design of the *in vitro* infection in RAW 264.7 cells. (B) Log CFU/mL of the intracellular bacteria at 2 and 18 h post infection. (C) TNF-α and (D) IL-6 levels in the cell free supernatant at 2 and 18 h post infection. Data are represented as mean ± SEM from three independent experiments (n = 3). Two-way ANOVA was performed to test the statistical significance, where *p<0.05, **p<0.01.

### Parthenolide improves survival and reduces systemic cytokine levels during oral infection

As Parthenolide proved to be a better anti-inflammatory molecule, we performed further *in vivo* infection experiments in mice (Figure 8A). We quantified bacterial burden in the liver, lung, and spleen after 2 days of treatment. Parthenolide treatment significantly reduced infection-induced systemic IL-6 and TNF-α production, while having little or no effect on systemic bacterial burden (Figure 8 B, C) It also increased the survival of treated mice significantly as compared to vehicle controls, indicating a protective effect during systemic infection (Figure 8 D). These findings remained consistent with the results from the *in vitro* infection experiments. Together, they suggested that Parthenolide did not have a direct role in curbing infection by reducing bacterial burden but rather protected the host reducing the systemic cytokine storm and eventually delaying the infection symptoms.

**FIGURE 8.**
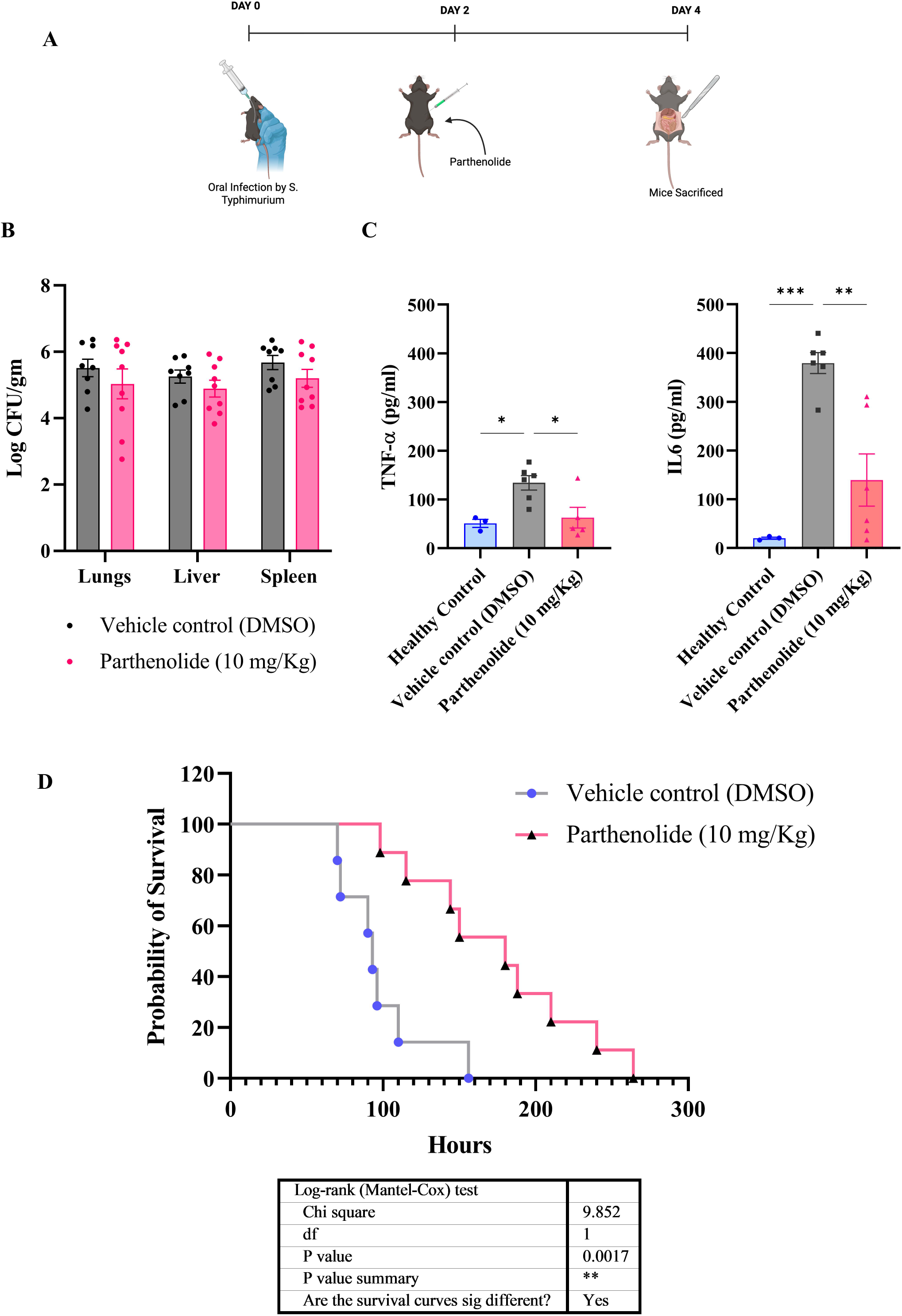
Parthenolide lowers TNF-α and IL-6 production and increases survival in orally infected C57BL/6 mice with *S*. Typhimurium. (A) Experimental design of oral infection in mice with *S*. Typhimurium. Post 2 days of infection, mice were injected intraperitoneally with Parthenolide (10 mg/kg) or DMSO (vehicle control). (B) Bacterial burden of vehicle control and Parthenolide treated mice, across different organs, lungs, liver and spleen. (C) Serum TNF-α and IL-6 levels in healthy control, vehicle control and Parthenolide treated mice. (D) Kaplan-Meier survival analysis comparing infected mice treated with DMSO or Parthenolide. Data are represented as mean ± SEM from three independent experiments (n = 3). Two-way ANOVA and one way ANOVA were performed to test the statistical significance, where *p<0.05, **p<0.01, ***p<0.001.

## DISCUSSION

Artemisinin and Parthenolide have been independently studied for their immunomodulatory effects. Popularly known for its use as an anti-malarial drug, Artemisinin and its derivatives have shown immunomodulatory activity *in vitro* and *in vivo* (16,17). Parthenolide, a traditional medicine used for migraine and arthritis (18), shows anti-cancer and immunomodulatory effects (19). However, direct comparisons between them under similar conditions have not been studied. Existing reports scattered across different cell types, disease models, and readouts often produce conflicting conclusions about each compound’s net effect on immune activation. In this study, we investigated the immunomodulatory effects of two sesquiterpene lactones, Artemisinin and Parthenolide on T cell and macrophage activation. Across different inflammatory conditions, Parthenolide behaved consistently as an anti-inflammatory compound, while the effects of Artemisinin were modest.

Parthenolide, in a dose-dependent manner suppressed cell blasting, IL-2 secretion and cell cycling upon activation (Figure 1, 2). It led to an increase in hypodiploid cells indicating impaired activation-associated proliferation, and this was mirrored by a reduction in both CD69 and CD44 activation markers (Figure 3). Artemisinin, tested under the same T cell activation conditions produced only a mild effect across these readouts. This pattern is consistent with earlier work showing that sesquiterpene lactones, including Parthenolide, suppress IL-2 expression and CD69 upregulation in stimulated blood T lymphocytes at high concentrations by blocking not only NF-κB but also NFAT and AP-1 DNA binding, transcription factors that act in parallel to drive the early T-cell activation program (20). In activated T cells, Parthenolide decreased the intracellular ROS levels (Figure 4). ROS generated downstream of TCR engagement, via NADPH oxidase 2 and mitochondrial electron transport, act as physiological second messengers required for full activation of NF- κB, AP-1, and downstream IL-2 production (21,22). These results are consistent with Parthenolide acting upstream, on signaling events that both activation and ROS production depend on. Artemisinin, by contrast, increased ROS in the same activated T cells.

Artemisinin and its derivatives have shown to suppress ROS levels in immune cells by activating *Nrf2* to provide antioxidant protection (23). This discrepancy could be attributed to the presence of the endoperoxide bridge of Artemisinin that mediates ROS generation bidirectionally (24).

To assess the effects of these compounds on other immune cells, we used primary macrophage population activated with LPS. Parthenolide inhibited LPS-induced nitrite levels substantially, while Artemisinin produced only a modest reduction by comparison. Parthenolide also lowered ROS levels more effectively than Artemisinin did, reinforcing Parthenolide’s ability to lower ROS and act as an anti-inflammatory compound post LPS activation. Cytokine measurements showed a more selective effect, Parthenolide reduced IL- 6 but had no effect on TNF-α, while Artemisinin produced only a slight reduction in IL-6 and TNF-α was unchanged. Neither compound altered overall metabolic activity in these primary cells (Figure 5). As adherent peritoneal macrophages are largely non-proliferative, we then turned to the RAW 264.7 macrophages to examine whether these effects extended to a proliferative macrophage population activated with LPS and IFN-γ, alone and in combination. Here again, Parthenolide again suppressed nitrite levels, but its cytokine profile slightly shifted. Unlike in primary macrophages, Parthenolide now suppressed both IL-6 and TNF-α slightly. Artemisinin’s profile also shifted, lowering nitrite while increasing IL-6 and modestly raising TNF-α, a divergence from its milder, largely neutral effect on cytokines in primary macrophages (Figure 6). Artemisinin’s ability to lower nitric oxide while simultaneously amplifying IL-6 and TNF-α suggests it may be acting on *Nos2* more directly than on the broader inflammatory cytokine program (25), whereas Parthenolide’s consistent suppression across nitrite, IL-6, and TNF-α points to a broader anti-inflammatory effect.

In the *in vitro* infection model, neither compound reduced intracellular *S.* Typhimurium burden in RAW 264.7 cells (Figure 7). Parthenolide likewise failed to reduce bacterial load in the liver, lung, or spleen of orally infected mice. Yet Parthenolide-treated mice survived significantly longer than controls, and this benefit was accompanied by reduced systemic IL- 6 and TNF-α rather than by any gain in bacterial clearance (Figure 8). This indicates that its benefit comes from limiting the host’s own inflammatory response rather than from any antimicrobial effect. This pattern is a hallmark of disease tolerance, a host defence strategy distinct from resistance, in which survival during infection is improved by limiting the damage caused by the immune response itself rather than by reducing pathogen burden (26).

Parthenolide can be positioned as a potential adjunct therapy specifically for infections such as typhoid fever or sepsis, where much of the tissue damage results from an overactive cytokine response (9,27,28), since a tolerance-based benefit does not extend to controlling the infection itself.

Overall, these findings position Parthenolide as a more promising candidate than Artemisinin for further development as a plant-derived immunomodulator (Figure 9). Structurally related sesquiterpene lactones acting on a shared signalling node can produce substantially divergent physiological consequences depending on cell type and activation state. In future, we will need to uncover the mechanisms by which Parthenolide acts as a better anti-inflammatory compound. Future work should test whether Parthenolide’s anti-inflammatory effects extend to other inflammatory diseases and systemic bacterial infections in which cytokine-driven pathology is the primary driver of host damage.

**FIGURE 9.**
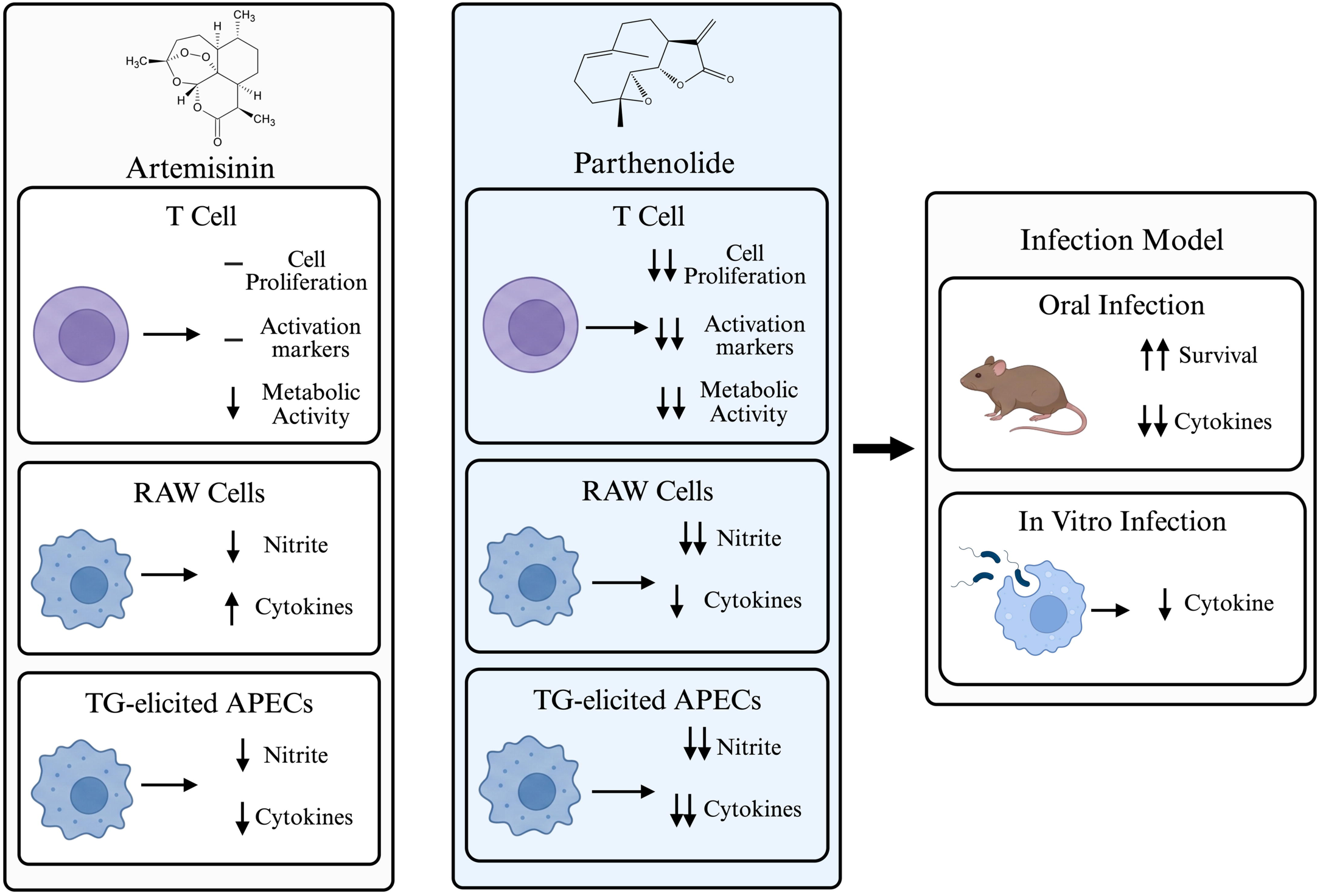
Comparative immuno-modulatory effects of Artemisinin and Parthenolide on immune cells and *S*. Typhimurium infection models. The differential effects of Artemisinin and Parthenolide on immune cell activation with respect to T cells, RAW 264.7 macrophage cell line and TG-elicited macrophages. In addition, the effects of these compounds on *S*. Typhimurium infection are represented. Arrows denote relative changes and double arrows indicate stronger effects.

## AUTHOR CONTRIBUTIONS

Tanisha Kumar: Methodology, Investigation, Formal analysis, Writing – original draft, review and editing.

Aagosh Kishor Karhale: Methodology, Investigation. Writing – review & editing. Shreyasee Das: Investigation, Methodology, Formal analysis, Writing – review & editing. Joel P Joseph: Investigation, Methodology, Supervision, Writing – reviewing & editing.

Dipankar Nandi: Conceptualization, Supervision, Writing – review & editing, Project administration, Funding acquisition.

## ACKNOWLEDGEMENTS

This work was supported by core grants from the Indian Institute of Science (IISc) and the DBT-IISc partnership grant. We are thankful for the infrastructural support from the FIST program of the Department of Science and Technology, India. We thank the staff of the Central Animal Facility, IISc, and the Flow Cytometry Facility, Division of Biological Sciences, IISc, for their support. We are grateful to all members of the DpN laboratory, especially Mabel D’Souza and Sai Rama Krishna Rongali, for their assistance with some experiments. The authors of this study have no conflict of interest to declare.

## REFERENCES

1. Sangeetha Vijayan P, Xavier J, Valappil MP. A review of immune modulators and immunotherapy in infectious diseases. Mol Cell Biochem. 2024 Aug;479(8):1937–55. doi:10.1007/s11010-023-04825-w

2. Strzelec M, Detka J, Mieszczak P, Sobocińska MK, Majka M. Immunomodulation—a general review of the current state-of-the-art and new therapeutic strategies for targeting the immune system. Front Immunol. 2023 Mar 9;14:1127704. doi:10.3389/fimmu.2023.1127704

3. Balasubramaniam M, Sapuan S, Hashim IF, Ismail NI, Yaakop AS, Kamaruzaman NA, et al. The properties and mechanism of action of plant immunomodulators in regulation of immune response – A narrative review focusing on Curcuma longa L., Panax ginseng C. A. Meyer and Moringa oleifera Lam. Heliyon. 2024 Apr;10(7):e28261. doi:10.1016/j.heliyon.2024.e28261

4. Shoaib M, Shah I, Ali N, Adhikari A, Tahir MN, Shah SWA, et al. Sesquiterpene lactone! a promising antioxidant, anticancer and moderate antinociceptive agent from Artemisia macrocephala jacquem. BMC Complement Altern Med. 2017 Dec;17(1):27. doi:10.1186/s12906-016-1517-y

5. Paço A, Brás T, Santos JO, Sampaio P, Gomes AC, Duarte MF. Anti-Inflammatory and Immunoregulatory Action of Sesquiterpene Lactones. Molecules. 2022 Feb 8;27(3):1142. doi:10.3390/molecules27031142

6. Pathak S, Gokhroo A, Kumar Dubey A, Majumdar S, Gupta S, Almeida A, et al. 7- Hydroxy Frullanolide, a sesquiterpene lactone, increases intracellular calcium amounts, lowers CD4+ T cell and macrophage responses, and ameliorates DSS-induced colitis. Int Immunopharmacol. 2021 Aug;97:107655. doi:10.1016/j.intimp.2021.107655

7. Tu Y. The discovery of artemisinin (qinghaosu) and gifts from Chinese medicine. Nat Med. 2011 Oct;17(10):1217–20. doi:10.1038/nm.2471

8. Qiu F, Liu J, Mo X, Liu H, Chen Y, Dai Z. Immunoregulation by Artemisinin and Its Derivatives: A New Role for Old Antimalarial Drugs. Front Immunol. 2021 Sep 9;12:751772. doi:10.3389/fimmu.2021.751772

9. He X, Wang C, Zhang R, Wang Y, Zhang Y, Yang T, et al. Parthenolide ameliorates inflammation in sepsis via covalently targeting Trim33 and inhibiting NF-κB pathway. Phytomedicine. 2026 Apr;153:157862. doi:10.1016/j.phymed.2026.157862

10. Aldieri E, Atragene D, Bergandi L, Riganti C, Costamagna C, Bosia A, et al. Artemisinin inhibits inducible nitric oxide synthase and nuclear factor NF kB activation. FEBS Lett. 2003 Sep 25;552(2–3):141–4. doi:10.1016/S0014-5793(03)00905-0

11. Saadane A, Masters S, DiDonato J, Li J, Berger M. Parthenolide Inhibits IκB Kinase, NF- κB Activation, and Inflammatory Response in Cystic Fibrosis Cells and Mice. Am J Respir Cell Mol Biol. 2007 Jun 1;36(6):728–36. doi:10.1165/rcmb.2006-0323OC

12. Joseph JP, Kumar T, Ramteke NS, Chatterjee K, Nandi D. High intracellular calcium amounts inhibit activation-induced proliferation of mouse T cells: Tert-butyl hydroquinone as an additive enhancer of intracellular calcium. Int Immunopharmacol. 2024 Dec;143:113501. doi:10.1016/j.intimp.2024.113501

13. Joseph JP, Gugulothu SB, Nandi D, Chatterjee K. Mechanical Properties Affect Primary T Cell Activation in 3D Bioprinted Hydrogels. ACS Macro Lett. 2023 Aug 15;12(8):1085–93. doi:10.1021/acsmacrolett.3c00271

14. Chattopadhyay A, Joseph JP, Jagdish S, Chaudhuri S, Ramteke NS, Karhale AK, et al. High throughput screening identifies auranofin and pentamidine as potent compounds that lower IFN-γ-induced Nitric Oxide and inflammatory responses in mice: DSS-induced colitis and Salmonella Typhimurium-induced sepsis. Int Immunopharmacol. 2023 Sep;122:110569. doi:10.1016/j.intimp.2023.110569

15. Chakraborty S, Banerjee P, Joseph JP, Pathak S, Verma T, Karhale AK, et al. Functional loss of rffG and rfbB, encoding dTDP-glucose 4,6-dehydratase, alters colony morphology, cell shape, motility and virulence in Salmonella Typhimurium. Front Microbiol. 2025 May 21;16:1572117. doi:10.3389/fmicb.2025.1572117

16. Li T, Chen H, Wei N, Mei X, Zhang S, Liu D lin, et al. Anti-inflammatory and immunomodulatory mechanisms of artemisinin on contact hypersensitivity. Int Immunopharmacol. 2012 Jan;12(1):144–50. doi:10.1016/j.intimp.2011.11.004

17. Long Z, Xiang W, Xiao W, Min Y, Qu F, Zhang B, et al. Advances in the study of artemisinin and its derivatives for the treatment of rheumatic skeletal disorders, autoimmune inflammatory diseases, and autoimmune disorders: a comprehensive review. Front Immunol. 2024 Oct 25;15:1432625. doi:10.3389/fimmu.2024.1432625

18. Wu C, Chen F, Rushing JW, Wang X, Kim HJ, Huang G, et al. Antiproliferative Activities of Parthenolide and Golden Feverfew Extract Against Three Human Cancer Cell Lines. J Med Food. 2006 Mar;9(1):55–61. doi:10.1089/jmf.2006.9.55

19. Mathema VB, Koh YS, Thakuri BC, Sillanpää M. Parthenolide, a Sesquiterpene Lactone, Expresses Multiple Anti-cancer and Anti-inflammatory Activities. Inflammation. 2012 Apr;35(2):560–5. doi:10.1007/s10753-011-9346-0

20. Humar M, Garcı a-Piñeres AJ, Castro V, Merfort I. Effect of sesquiterpene lactones on the expression of the activation marker CD69 and of IL-2 in T-lymphocytes in whole blood. Biochem Pharmacol. 2003 May;65(9):1551–63. doi:10.1016/S0006-2952(03)00108-4

21. Franchina DG, Dostert C, Brenner D. Reactive Oxygen Species: Involvement in T Cell Signaling and Metabolism. Trends Immunol. 2018 Jun;39(6):489–502. doi:10.1016/j.it.2018.01.005

22. Kesarwani P, Murali AK, Al-Khami AA, Mehrotra S. Redox Regulation of T-Cell Function: From Molecular Mechanisms to Significance in Human Health and Disease. Antioxid Redox Signal. 2013 Apr;18(12):1497–534. doi:10.1089/ars.2011.4073

23. Hua L, Liang S, Zhou Y, Wu X, Cai H, Liu Z, et al. Artemisinin-derived artemisitene blocks ROS-mediated NLRP3 inflammasome and alleviates ulcerative colitis. Int Immunopharmacol. 2022 Dec;113:109431. doi:10.1016/j.intimp.2022.109431

24. Zhou F, Li G, Tan R, Wu G, Deng C. Artemisinins in autoimmune diseases: effects and mechanisms in systemic lupus erythematosus and rheumatoid arthritis. Br J Pharmacol. 2025 Aug;182(15):3411–27. doi:10.1111/bph.70086

25. Yuan DS, Chen YP, Tan LL, Huang SQ, Li CQ, Wang Q, et al. Artemisinin: A Panacea Eligible for Unrestrictive Use? Front Pharmacol. 2017 Oct 17;8:737. doi:10.3389/fphar.2017.00737

26. Medzhitov R, Schneider DS, Soares MP. Disease Tolerance as a Defense Strategy. Science. 2012 Feb 24;335(6071):936–41. doi:10.1126/science.1214935

27. Sheehan M, Wong HR, Hake PW, Zingarelli B. Parthenolide improves systemic hemodynamics and decreases tissue leukosequestration in rats with polymicrobial sepsis*: Crit Care Med. 2003 Sep;31(9):2263–70. doi:10.1097/01.CCM.0000085186.14867.F7

28. Everest P, Wain J, Roberts M, Rook G, Dougan G. The molecular mechanisms of severe typhoid fever. Trends Microbiol. 2001 Jul;9(7):316–20. doi:10.1016/S0966-842X(01)02067-4

