## Supplemental Information for "Comparative analysis of immunomodulatory effects of Artemisinin and Parthenolide on activation of immune cells and their roles in *Salmonella* Typhimurium infection"

**SUPPLEMENTARY FIGURES**

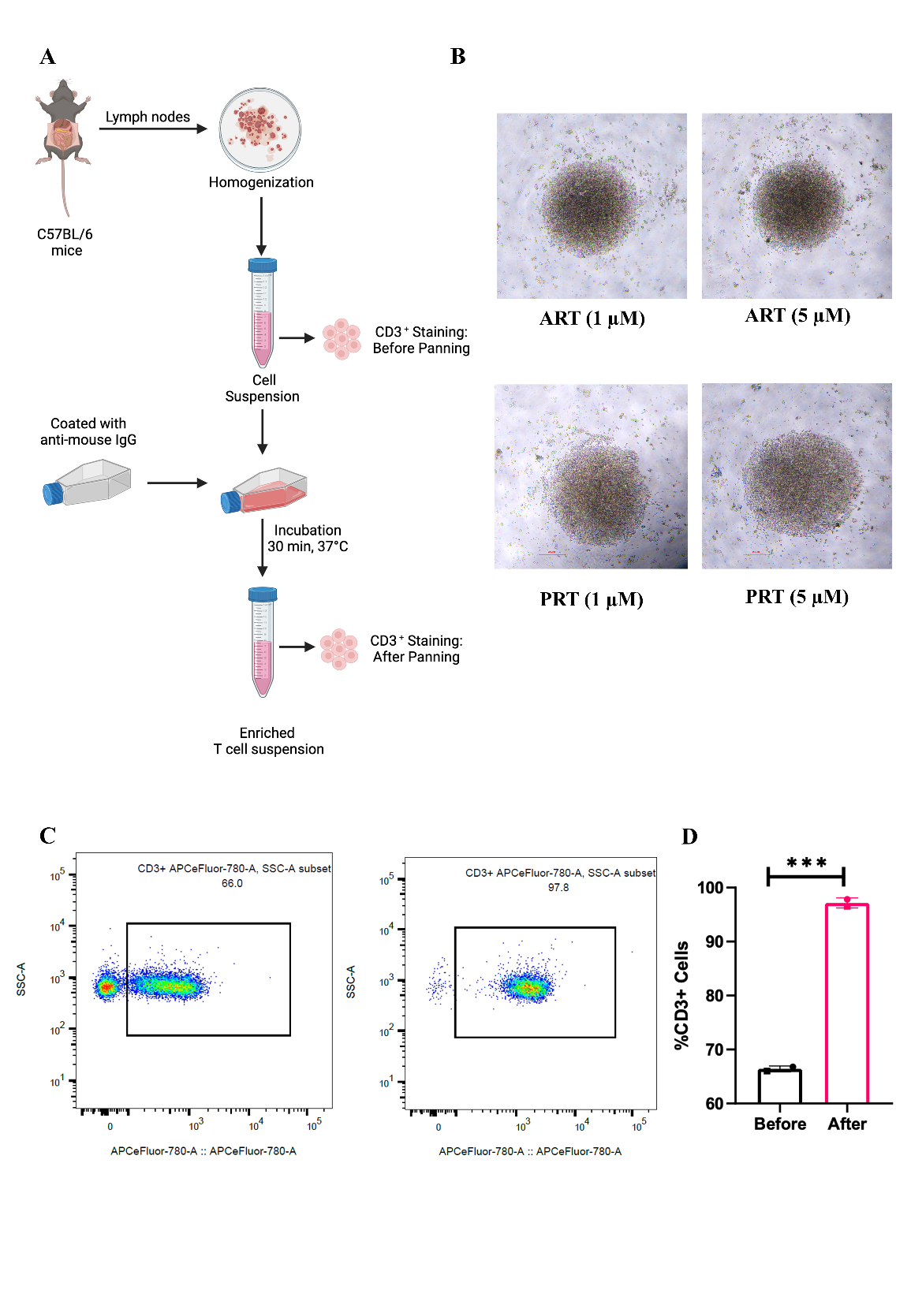

SUPPLEMENTARY FIGURE 1. Established protocol gives CD3^+^ enriched cell suspension.
(A) Schematic representation of the experimental flow to obtain CD3^+^ enriched cell suspension. (B) Flow cytometric plots of CD3^+^ cells before and after panning. (C) Quantification of flow cytometry data. (D) Brightfield images of unactivated T cells upon Artemisinin and Parthenolide treatment. Data are represented as mean ± SEM from two independent experiments. Unpaired t-test was performed to test the statistical significance, where ***p<0.001.

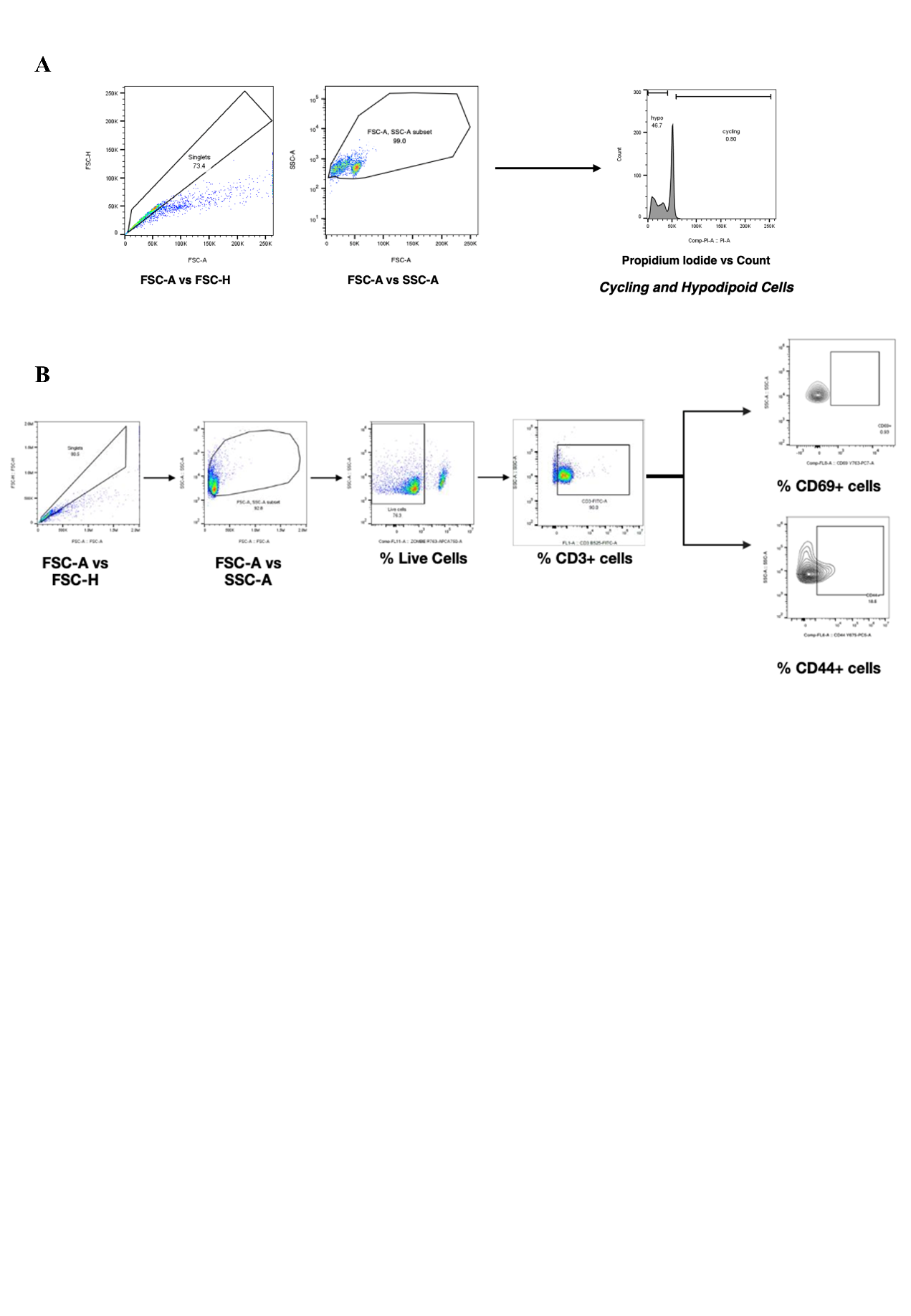

SUPPLEMENTARY FIGURE 2. Representative gating strategies for flow cytometry experiments performed to determine (A) percentage of cycling and hypodiploid cells and (B) percentage of CD69^+^ and CD44^+^ cells.

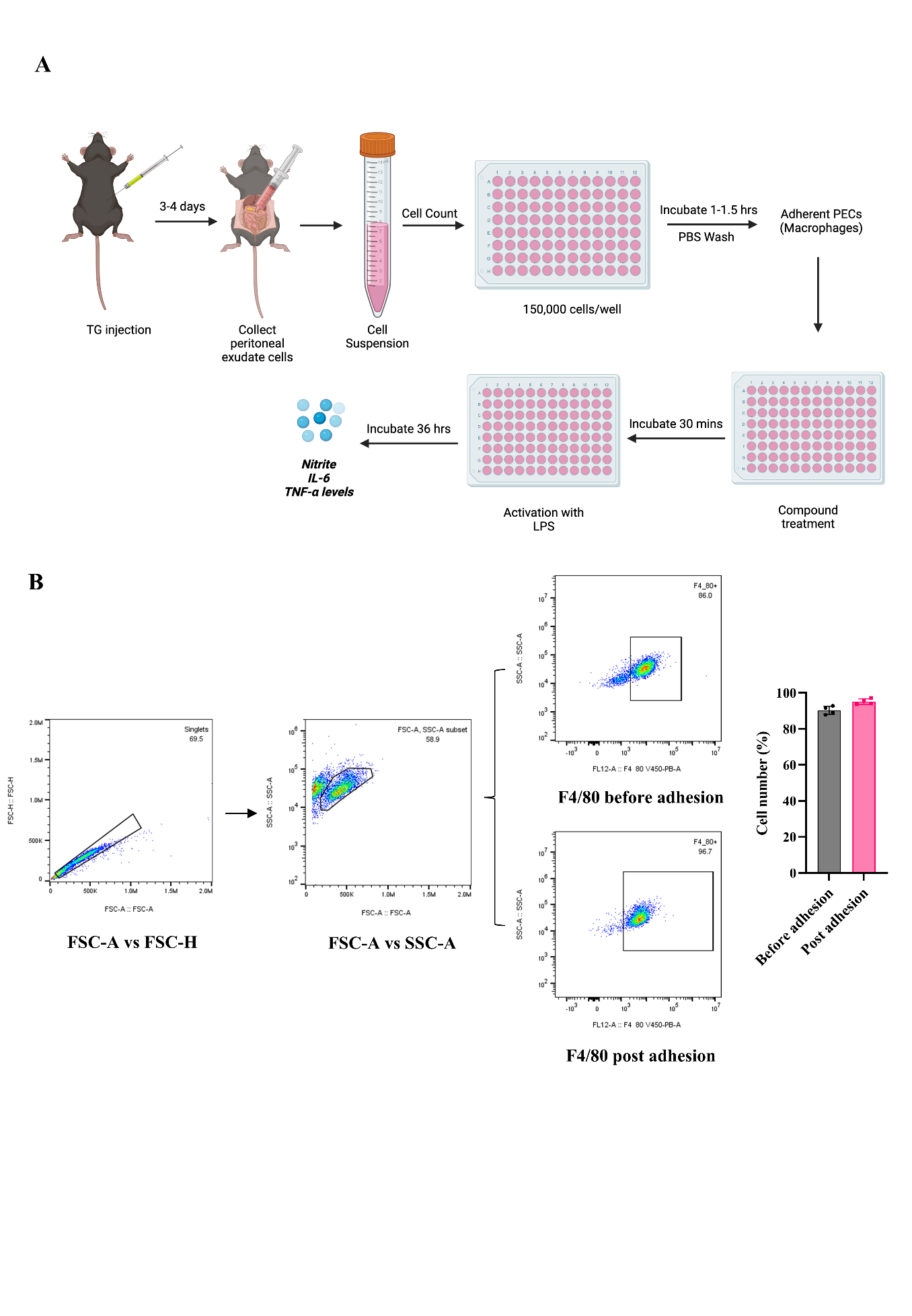

SUPPLEMENTARY FIGURE 3. Thioglycolate- elicited peritoneal exudate cells are enriched in F4/80+ macrophages. (A) Schematic representation of the experimental flow of the extraction of TG-elicited APECs and treatment with Artemisinin and Parthenolide and activation with LPS. (B) Flow cytometric plots of F4/80+ cells before and after adhesion and quantification of the flow cytometry data. Data are represented as mean ± SEM from four independent experiments.

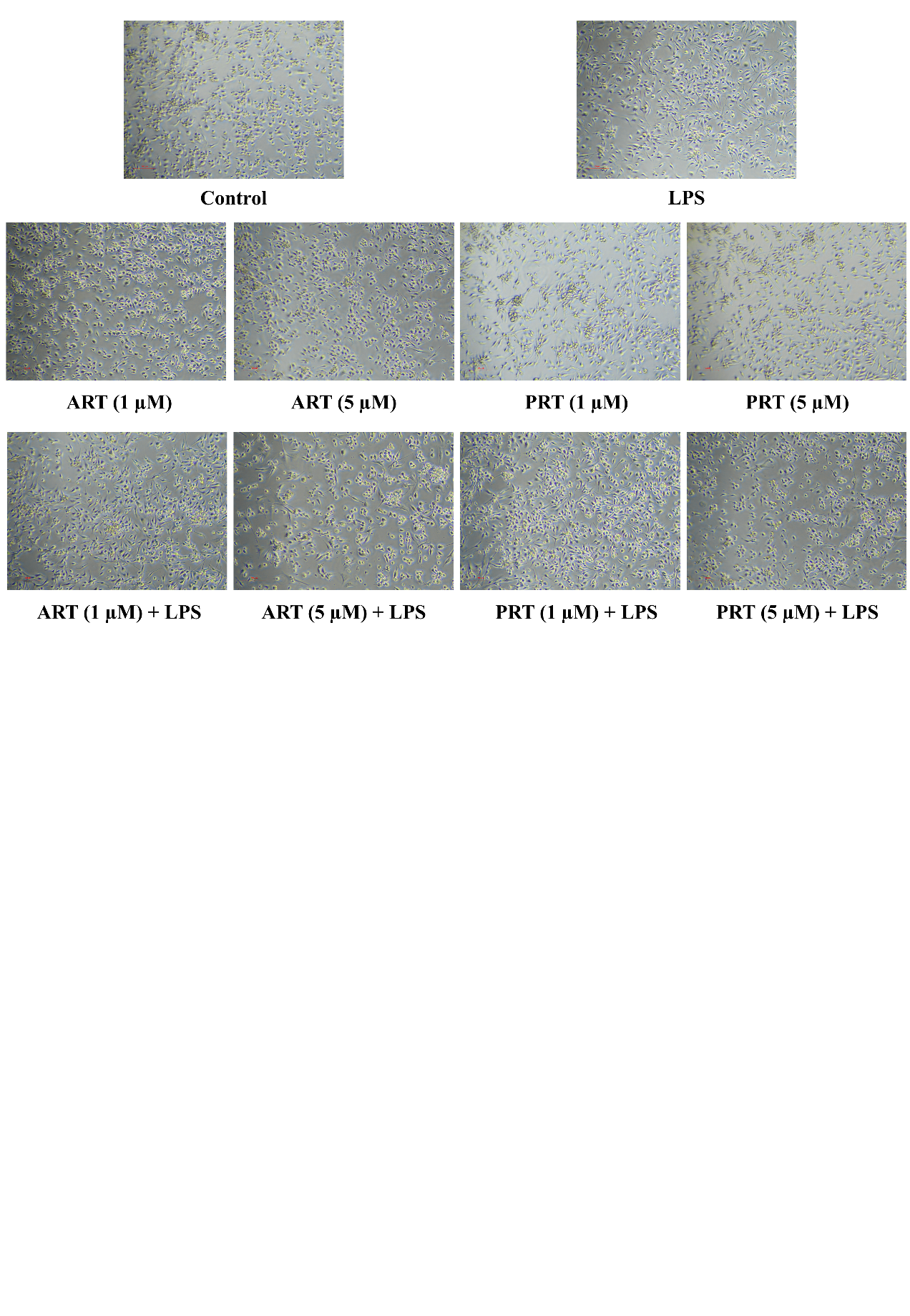

SUPPLEMENTARY FIGURE 4. Brightfield images of TG-elicited peritoneal macrophages activated with LPS and treated with Artemisinin and Parthenolide at different concentrations.

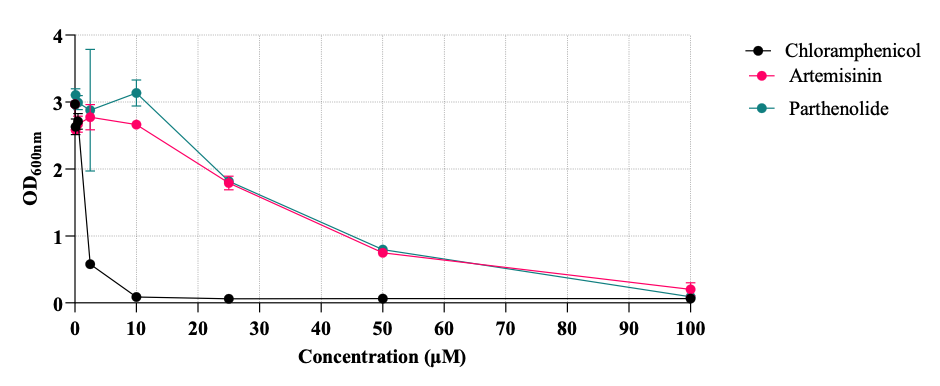

SUPPLEMENTARY FIGURE 5. Growth of *S*. Typhimurium in LB is not affected by Parthenolide or Artemisinin treatment but is greatly reduced by Chloramphenicol, an antibiotic.

**SUPPLEMENTARY METHODS**

### **Cell purity test**

The percentage purity of T cell suspension and adherent macrophages was determined by staining the cells for its surface markers. The cells were incubated with pre-titrated dilutions of anti-mouse CD3 at 1:100 of the conjugated antibodies for T cells and F4/80 for macrophages. The cells were incubated in the dark on ice for 45 minutes with intermittent tapping. The cells were washed with FACS Buffer (PBS + 2% FBS + 0.1% Sodium Azide) and fixed with 1% paraformaldehyde in PBS.

**Compound efficacy on growth of *Salmonella* Typhimurium**

Briefly, an overnight primary culture of *Salmonella* Typhimurium 14028s was grown in Luria-Bertani (LB) media at 37°C with orbital shaking at 160 rpm. A secondary culture was prepared to an absorbance of 1.0, of which 0.2% (v/v) was inoculated into fresh LB media supplemented with varying concentrations of Artemisinin or Parthenolide, as indicated in the respective figures. Chloramphenicol served as the positive control. All cultures were incubated at 37°C with orbital shaking at 160 rpm. Following six hours of incubation, the absorbance was measured at 600 nm using a TECAN microplate reader.
